# Dietary restriction and life-history trade-offs: insights into mTOR pathway regulation and reproductive investment in Japanese quails

**DOI:** 10.1101/2023.11.14.567012

**Authors:** Gebrehaweria Kidane Reda, Sawadi Fransisco Ndunguru, Brigitta Csernus, Gabriella Gulyás, Renáta Knop, Csaba Szabó, Levente Czeglédi, Ádám Z. Lendvai

## Abstract

Resources are needed for growth, reproduction and survival, and organisms must trade-off limited resources among competing processes. Nutritional availability in organisms is sensed and monitored by nutrient-sensing pathways that can trigger physiological changes or alter gene expression. Previous studies have proposed that one such signalling pathway, the mechanistic target of rapamycin (mTOR), underpins a form of adaptive plasticity when individuals encounter constraints in their energy budget. Despite the fundamental importance of this process in evolutionary biology, how nutritional limitation is regulated through the expression of genes governing this pathway and its consequential effects on fitness remains understudied, particularly in birds. We used dietary restriction to simulate resource depletion and examined its effects on body mass, reproduction and gene expression in Japanese quails (*Coturnix japonica*). Quails were subjected to *ad libitum* (ADL) feeding or 20%, 30%, and 40% restriction levels for two weeks. All restricted groups exhibited reduced body mass, whereas reductions in the number and mass of eggs were observed only under more severe restrictions. Dietary restriction led to decreased expression of *mTOR* and insulin-like growth factor 1 (*IGF1*), whereas the ribosomal protein S6 kinase 1 (*RPS6K1*) and autophagy-related genes (*ATG9A* and *ATG5*) were upregulated. The pattern in which mTOR respond to restriction was similar to what has been seen in body mass. Regardless of the treatment, proportionally higher reproductive investment was associated with individual variation in *mTOR* expression. These findings reveal the connection between dietary intake and the expression of *mTOR* and related genes in this pathway.

## 1. INTRODUCTION

Resource availability is a key driver of resource allocation decisions which define life-history trade-offs (Ng’oma et al., 2017; Zera and Harshman, 2001). When food is abundant, animals allocate resources towards current reproduction against somatic maintenance and further reproduction. The bias for reproductive investment may be accompanied by physiological costs, including oxidative stress and a reduced immune potential (Chang van Oordt et al., 2022; Metcalfe and Monaghan, 2013), which in turn affects future reproduction performance and the health span of the organism (Hassan et al., 2003; Mahrose et al., 2022). This phenomenon changes under limited resources when organisms must divert energy from reproduction to somatic maintenance (Carlsson et al., 2021; Flatt et al., 2013).

Dietary restriction (DR) is an intervention that mimics the depletion of resources, which regulates life-history traits almost uniformly in model organisms ranging from yeast to humans (Colman et al., 2014; Inness and Metcalfe, 2008; Simons et al., 2013). Reducing calorie intake affects the investment into self-maintenance, growth and reproduction antagonistically (McCracken et al., 2020; Regan et al., 2020). The classical resource allocation theory predicts that DR should lead to a linear decrease in reproduction in favour of self-maintenance (Shanley and Kirkwood, 2000). However, modest DR can maintain or even improve reproductive performance by activating cell recycling mechanisms such as apoptosis and autophagy (Adler and Bonduriansky, 2014; Mahrose et al., 2022). When the level of DR becomes more severe, the organism must shift energy away to meet basal energetic requirements; thus, the reproduction rate will decline (Moatt et al., 2016; Ottinger et al., 2005; Shanley and Kirkwood, 2000).

Resource availability at the organismal level is monitored by the neuroendocrine system that dynamically responds to internal signals via changes in physiology or gene expression (Maruska et al., 2018). The key mediators of this process are the nutrient-sensing pathways governed by insulin-like growth factor-1 (IGF-1) and mechanistic target of rapamycin (mTOR) (Johnson, 2018; Kapahi et al., 2017). The IGF-1/mTOR signalling pathway is activated by high nutrient availability and triggers growth and reproduction while downregulating cellular processes that maintain organismal and cellular homeostasis (e.g. apoptosis and autophagy) (Montoya et al., 2022; Papadopoli et al., 2019). In response to growth hormone and energy availability, IGF-1 is released into the bloodstream and binds to its membrane receptor (IGF-1R), which activates further molecular components (PI3K and Akt) that will, in turn, trigger mTOR activation (Feng and Levine, 2010).

mTOR serves as the central regulator of the nutrient-sensing pathway, integrating intracellular nutrient availability and extracellular signals to govern essential cellular processes, including metabolism, growth, proliferation, and survival, thereby influencing tissue and organ growth (Papadopoli et al., 2019; Rabanal-Ruiz and Korolchuk, 2018). The mTOR exists in two distinct complexes known as mTORC1 and mTORC2, each with different functions and regulatory mechanisms. mTORC1 is primarily involved in regulating cell growth and protein synthesis in response to various environmental cues such as nutrient availability, energy status, and growth factors. On the other hand, mTORC2 has diverse roles in cell survival, cytoskeletal organization, and metabolism (Szwed et al., 2021).

DR has been shown to downregulate the mTORC1 pathway through various mechanisms, primarily involving its upstream effectors, such as IGF-1 and intracellular amino acid deprivation (Sancak et al., 2010; Speakman and Mitchell, 2011). Furthermore, DR has a complex impact on mTORC2 activity. On the one hand, it has been observed that DR downregulates mTORC2 by inhibiting insulin/Akt signalling (Saxton and Sabatini, 2017). On the other hand, there is evidence to suggest that DR can upregulate mTORC2 through the activation of adenosine monophosphate-activated protein kinase (AMPK) (Fu and Hall, 2020).

The activity of mTOR and its downstream effectors is further regulated through transcriptional regulation/mRNA expression and post-translational modifications (Deng et al., 2014; Mierziak et al., 2021; Rollins et al., 2019). While post-translational modifications and specific amino acid availability induce mTOR activation, general resource availability is responsible for adaptive changes in gene expression (Efeyan et al., 2015; Mierziak et al., 2021; Sandri et al., 2013). Although the final activity of mTOR is influenced by multiple factors, higher expression of the mTOR gene can potentially increase the pool of available mTOR protein for activation, while lower gene expression leads to reduced protein production and availability (Buccitelli and Selbach, 2020). Studies on model organisms have predominantly focused on the post-translational activation of mTOR and its downstream effectors (Laplante and Sabatini, 2012; Papadopoli et al., 2019). However, the impact of DR on the differential expression of nutrient sensing genes and their role in mediating fitness remain to be fully elucidated.

The mTORC1 performs its effect on fitness traits through a number of downstream effectors, including ribosomal protein S6 kinase 1 (RPS6K1) (Guo and Yu, 2019; Nojima et al., 2003) and autophagy-related genes (Ma et al., 2018). Under excessive food availability, the mTORC1/RPS6K1 pathway facilitates protein synthesis and subsequent cell growth and proliferation (Fenton and Gout, 2011). Under DR, the mTORC1/autophagy pathway recycles cell contents for energy substitution and reduces oxidative stress (Chung and Chung, 2019). Hence, *RPS6K1*, *ATG9A* and *ATG5* are the best representative candidate genes of the mTOR downstream pathways. Studying the expression of these genes and their relationship to fitness traits under different DR levels is critically important to understand their role beyond post-translational regulation.

Previous studies on DR have primarily focused on model organisms other than birds, and except for some production-intentioned experiments on chicken (Deng et al., 2014; Hao et al., 2021; She et al., 2019), understanding the mechanism of mTOR signalling in avian biology remains largely unclear. Studies conducted in mammals have indicated that a higher nutritional intake and metabolic rate upregulate the mTOR pathway, leading to a subsequent downregulation of autophagy. This, in turn, exposes organisms to oxidative damage and cellular senescence. However, DR without malnutrition has been found to mitigate these effects (Carroll and Korolchuk, 2018). On the contrary, birds have higher metabolic rates, circulating glucose levels and body temperature than mammals, while they live twice as long as size-matched mammals (Barja, 1998). Despite the fact that a higher metabolic rate contributes to oxidative and glycoxidative damages, birds suffer less compared to mammals at a given body size (Jimenez et al., 2019). This metabolic paradox may be due to a different diet-fitness relationship in birds, which can be elucidated using dietary manipulation experiments.

At this stage, we aimed to observe the effects of DR gradient on liver *mTOR* mRNA expression, as well as its main upstream and downstream effectors, using adult female Japanese quails (*Coturnix japonica*) as an experimental avian model system. The liver serves as a primary site for the intricate nutrient metabolic pathway, which significantly influences the proper functioning of the entire body. Being a major organ responsible for regulating homeostasis, the liver plays a crucial role in nutrient regulation, protein synthesis, and detoxification processes. Nearly all genes involved in the nutrient-sensing pathway exhibit differential expression in the liver and display a robust correlation with the overall functioning of the body, ultimately determining fitness traits (Baloni et al., 2019; Gokarn et al., 2018).

We hypothesize that under DR, the expression of target genes involved in the mTOR pathway plays a crucial role in mediating resource availability and influencing fitness traits. We predict reduced *mTOR*, *IGF1* and *RPS6K1* expression and upregulation of autophagy-related genes. Furthermore, we anticipate that variations in *mTOR* expression will be associated with proportional differences in reproductive investment. These findings will provide insights into the relationship between gene expression, resource allocation, and fitness traits.

## 2. Materials and methods

### 2.1. Experimental animals and housing

The experiment was approved by the Ethical Committee for animal use of the University of Debrecen, Hungary (Protocol No. 5/2021/DEMAB) and followed all institutional and national regulations.

We purchased four weeks old Japanese quail (*Coturnix japonica*) chicks from a commercial quail breeder (Budai Fürjészet, Hungary) and housed them in the animal house of the Institute of Agricultural Research and Educational Farm of the University of Debrecen. The birds were kept in cages in groups of 10 for an additional four weeks (until they reached maturity) before being subjected to the experimental treatment. At the age of eight weeks, 32 female birds with similar body mass were selected and housed in individual cages (18.5 cm long × 21 cm wide × 18.5 cm high) for seven days of acclimation period on *ad libitum* (ADL) feed and water. The experimental room was maintained at a temperature of 24±3 °C and 60 - 75% relative humidity. Photoperiod was fixed at 12:12 L:D daily cycle and regulated using a LED Lighting Dimming System. The basal feed for experimental quails was formulated as a breeder quail ration (20% CP; 12.13 MJ/kg ME, (NRC (National Research Council), 1994)) based on corn, soybean, and wheat (Table S1).

### 2.2. Experimental design

During the acclimation period, the daily feed intake of each individual was measured for seven consecutive days. Approximately 50 g feed was weighed on a digital scale (± 0.1 g) and provided in a 200 g capacity plastic feeder each morning between 8:00-9:00 am. The following day, 24h later, the weight of the remaining feed was weighed again, and the feeders were replenished with fresh feed. Daily feed intake was measured as the difference between the weight of the offered and the remaining food. Due to the design of the feeders, food spillage was negligible. The average daily feed intake was calculated as the mean of the seven measurements. The average feed intake during the acclimation period was 29.67 ± 3.73 g. We also measured the live body mass of each bird at the beginning and at the end of the acclimation period to analyse mass change. At the start of the experimental treatment, there were no significant temporal pattern of either body mass or feed intake. Birds were regularly laying eggs during the acclimation period.

After acclimation, 32 female birds were randomly assigned to four dietary treatments. The birds were fed with 80% (DR20), 70% (DR30), and 60% (DR40) of their average individual feed intake, while the control group was fed ad libitum (ADL). The experiment lasted for 14 days. The amount of feed left daily in the ADL group was measured and analysed to detect any significant changes in temporal intake. However, no significant changes were observed.

### 2.3. Measurements and sampling

Immediately after lights-on in the morning, we removed all the feeders to maintain similar feeding conditions (empty gut) between the ADL and restricted birds during the measurement and sampling points. We measured body mass at the beginning of the experiment (day 0) and on days 7 and 14 of the restriction period. We collected eggs daily, measured their mass and recorded the identity of the respective hen. Body mass and egg mass were measured using a digital scale (± 0.1 g). On day 14, all birds were euthanised, and immediately a liver sample was collected and rapidly frozen on dry ice, and immediately stored at –80 ℃ until further assays.

### 2.4. RNA Extraction and cDNA Synthesis

Total RNA from liver tissue was isolated using the TRIzol reagent (Direct-zol™ RNA MiniPrep, Zymo Research Corporation, USA) according to the manufacturer’s protocol, including DNA digestion step. The RNA concentration and purity were measured using HTX Synergy Multi-Mode Microplate Reader spectrophotometer (Agilent BioTek, BioTek Instruments Inc, USA). RNA integrity was checked by 1% agarose gel electrophoresis stained with ethidium bromide (see supplementary material for more detailed protocol). Reverse transcription was performed using the qScript cDNA synthesis kit, following the manufacturer’s protocol (Quantabio Reagent Technologies, QIAGEN Beverly Inc., USA) in PCRmax Alpha Thermal Cycler (Cole-Parmer Ltd., Vernon Hills, IL, USA) (see supplementary material for more detailed protocol).

### 2.5. Real-time PCR (qPCR)

The real-time quantitative PCR was performed using EvaGreen qPCR Mix (*Solis BioDyne*, Teaduspargi, Estonia) according to the manufacturer’s protocol. Intron-spanning gene-specific primer pairs for quails were designed using Oligo7 software and obtained from Integrated DNA Technologies (BVBA-Leuven, Belgium) (Table S2). We checked for target identity using Primer-Blast software of the National Centre for Biotechnology Information (NCBI) (http://www.ncbi.nlm.nih.gov) (see Supplementary Material for more detailed protocol).

Among the most frequently used reference genes in birds, beta-actin (*ACTB*), glyceraldehyde-3-phosphate dehydrogenase (*GAPDH*) and 18S ribosomal RNA (RN18S), we selected the best reference gene, *ACTB*, using NormFinder, BestKeeper and deltaCt algorithms. The 2^−ΔΔCt^ method was employed to analyse the relative changes in mRNA expression of target genes (*mTOR*, *RPS6K1*, *IGF1*, *IGF1R, ATG9A* and *ATG5*) (Livak and Schmittgen, 2001).

### 2.6. Statistical analysis

All analyses were performed using R v. 4.1.2 ‘Bird Hippie’ (R Core Team, 2021). We fitted four models to analyse our data depending on the data source and relationship of variables. We used a linear model to analyse single-time data such as relative mRNA expression, hereafter gene expression, and the regression of one-to-one parameters. To analyse body mass across restriction time and restriction levels, we used linear mixed-effects models using ‘lme4’ (Bates et al., 2015) and ‘lmerTest’ packages v. 3.1-3 (Kuznetsova et al., 2017) considering individuals and experimental blocks as a random intercept (Table S3). To analyse the egg mass across days and restriction level, we used generalised additive mixed-effects models using the ‘mgcv’ package v. 1.8-40 (Wood, 2017) to incorporate nonlinear forms of the predictor restriction days. We used generalised linear mixed-effects models of the family logit using package ‘aod’ v. 1.3.2 (Lesnoff and Lancelot, 2012) to analyse binary response variable (daily egg laying). In mixed models, the individual bird ID was included as a random intercept to control for the repeated measures. Akaike’s information criterion (AICc) was used to choose the best-supported models (Burnham and Anderson, 2010). The log of fold change was used to analyse the relative gene expression. The Tukey test was used as *post hoc* test with p < 0.05 significance level and bars set as means ± SEM. To see the multivariate regression of genes’ mRNA expression against the fitness traits and resource allocation strategy, we used principal component analysis (PCA) using ‘prcomp’ function from ‘stats’ package to avoid multicollinearity between the predictor variables (R Core Team, 2021). We used the ‘ggbiplot’ package to visualise the clustered data against treatments (Vu, 2011). We used Kaiser’s rule to retain PCs for further analysis (Kaiser, 1960). We analysed individual genes’ impact on each fitness traits using linear regression. We analysed resource allocation strategy using the proportional change in egg mass in function of change in body mass compared to the pre-treatment status. We used trigonometric calculation to determine the direction of the resulting vector in radians (–π/2 - π/2) that reflects resource allocation strategy. A value of zero corresponds to no re-allocation, positive and negative values indicate re-allocation towards reproduction and self-maintenance, respectively. This method allowed us to analyse the relationship between individual gene expression and resource allocation strategy across treatment groups.

## 3. Results

### 3.1. Dietary restriction affects body mass

Experimental groups did not differ in their initial body mass, but DR reduced it significantly (treatment: *F_3,28_* = 13.832, *p* < 0.0001; time: *F_2,56_* = 61.826, *p* < 0.0001; treatment × time interaction: *F_6,56_* = 12.262, *p* < 0.0001). Birds grouped under all restriction levels showed a significant reduction in body mass compared to the *ad libitum*-fed birds (ADL) at both week 1 and week 2. The DR40 also resulted in significantly lower body mass than the DR20 at both time points, while the other restricted groups did not differ significantly (Figure 1, Table S4). All restricted groups showed significantly lower body mass at both weeks compared to initial body mass. The DR30 and DR40 groups showed a significant and marginally significant body mass reduction from week 1 to week 2, respectively, while the DR20 group showed no further significant variation from week 1 to week 2 restriction time points (Table S5).

**FIGURE 1.**
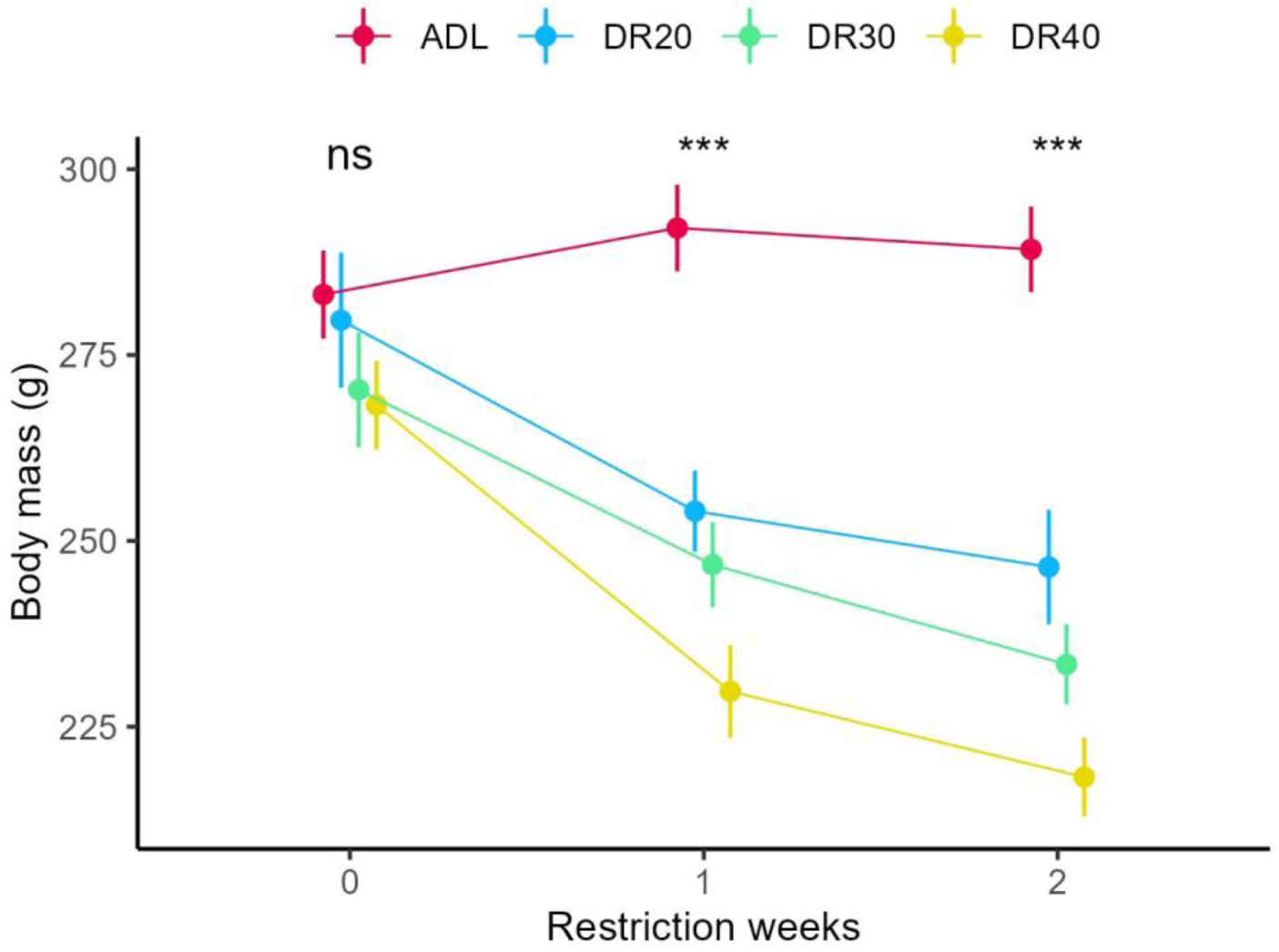
The effect of different dietary restriction levels on body mass of female quails at different time points. Data are represented by the mean ± SE from 8 birds per group. ns: none significant at p < 0.05; *** significantly different at p < 0.001; ADL: *ad libitum*; DR20: 20% restriction; DR30: 30% restriction; DR40: 40% restriction.

### 3.2. Severe dietary restriction reduces reproductive traits

The treatment groups differed significantly in the total number of eggs laid during the 14 days restriction period (*F_3,24_* = 5.448, *p* < 0.005). Restricted birds decreased daily egg-laying probability in compare to the ADL group. Additionally, overall probability of daily egg laying was significantly reduced across restriction period (Table 1, Figure 2a). Concerning total number of eggs, the DR40 group laid significantly fewer eggs than the ADL group (*t* = 3.992, *p* = 0.002), while the DR20 and DR30 did not show significant variation with the ADL (Figure 2b). The level of DR and restriction period significantly explained daily egg-laying probability.

**FIGURE 2.**
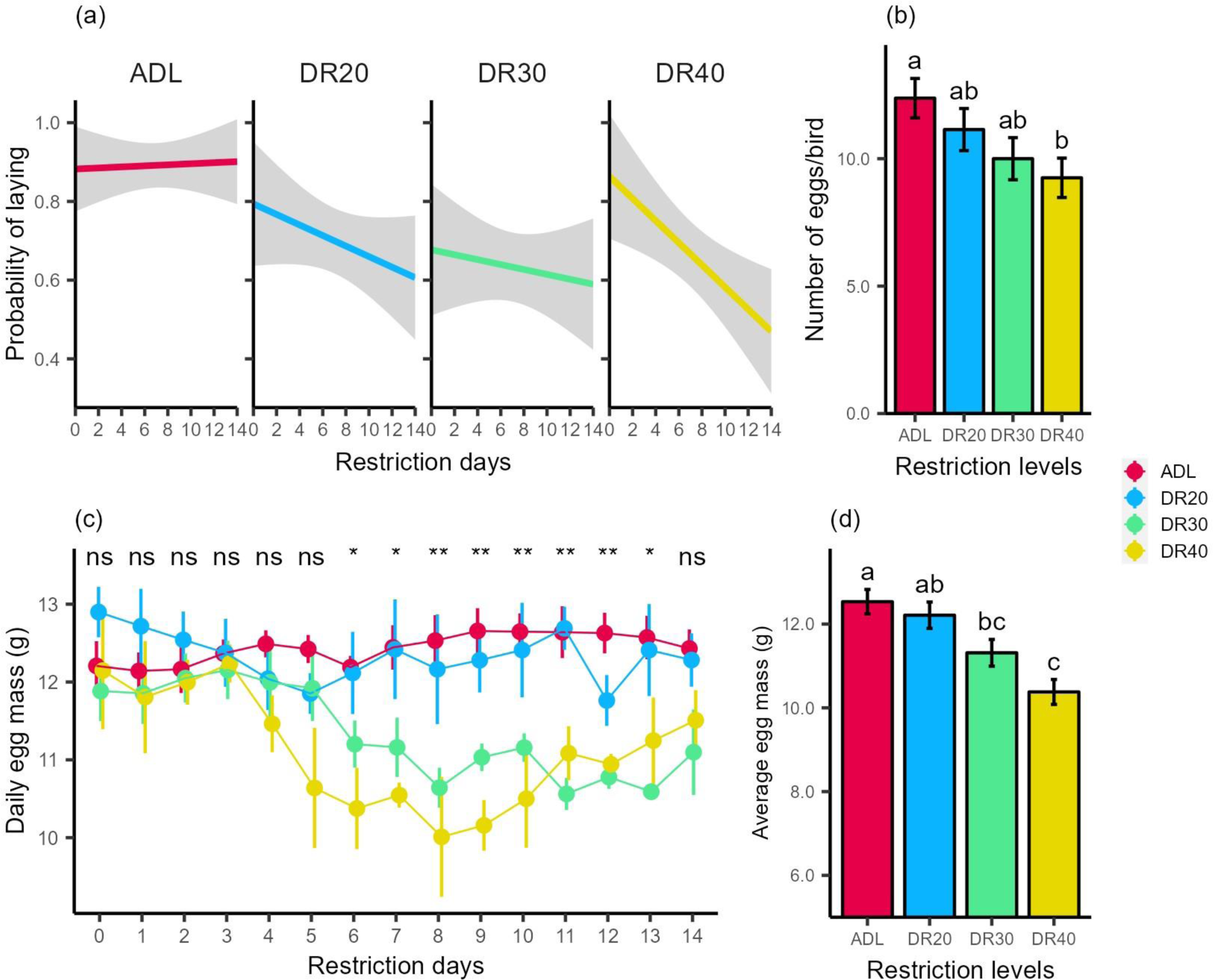
Effect of dietary restriction on egg production and egg mass. While all the restricted groups slightly reduce probability of daily egg laying, the DR40 showed the highest reduction (a). Only DR40 significantly reduced total number of eggs (b). Egg mass reduced at DR30 and DR40 starting the five sixth day (c). Total egg mass was significantly lower in the DR30 and DR40 groups. Data are represented by the mean ± SE from 8 birds per treatment group. Data are represented by the mean ± SE from 8 birds per group. Means followed by a common letter are not significantly different at p < 0.05. ns: none significant; * significantly different at p < 0.05; ** significantly different at p < 0.01. Abbreviations follow Fig. 1.

**TABLE 1.** Generalised linear mixed model of the family logit predicting the probability of daily egg laying as a function of dietary restriction levels restriction days with restriction treatment and restriction days as fixed effect and individual bird as random effect.

| *Predictors* | *Log-Odds* | *SE* | *z value* | *p-value* |
| --- | --- | --- | --- | --- |
| Intercept | 2.82 | 0.53 | 5.29 | 0.0001 |
| DR20 | -1.57 | 0.65 | -2.39 | 0.0154 |
| DR30 | -1.69 | 0.65 | -2.57 | 0.0089 |
| DR40 | -1.58 | 0.64 | -2.44 | 0.0132 |
| Day | -0.06 | 0.02 | -2.41 | 0.0151 |
| *Random effects* |  |  |  |  |
| birdID Variance | 1.127 |  |  |  |
| Observations | 480 |  |  |  |

birdID: individual bird, SE: standard error

The time-dependent trend indicated that egg mass was significantly reduced in DR30 and DR40 groups starting from day 5 (*p* < 0.003). Like the ADL group, egg mass from the DR20 group showed no change throughout the restriction period. On the last days of the experiment, egg mass from the DR30 and DR40 groups showed improvement to the level that they no longer differed from the ADL and DR20 groups (Figure 2c), Table S6). Average egg mass in the two weeks restriction period was significantly lower in the DR30 and DR40 groups compare to the ADL fed groups, while the DR40 group still showed significantly lowed average egg mass than the DR20 (Figure 2d).

### 3.3. Dietary restriction affects gene expression

The *mTOR* gene expression showed a significant and gradual decrease across the DR levels (*F_3,28_* = 15.424, *p* < 0.0001; Figure 3a, Table S7). However, the expression of *RPS6K1* was increased in response to the treatments (*F_3,28_* = 7.522, *p* < 0.001), showing an increasing trend with the severity of the restrictions (Figure 3b, Table S7).

**FIGURE 3.**
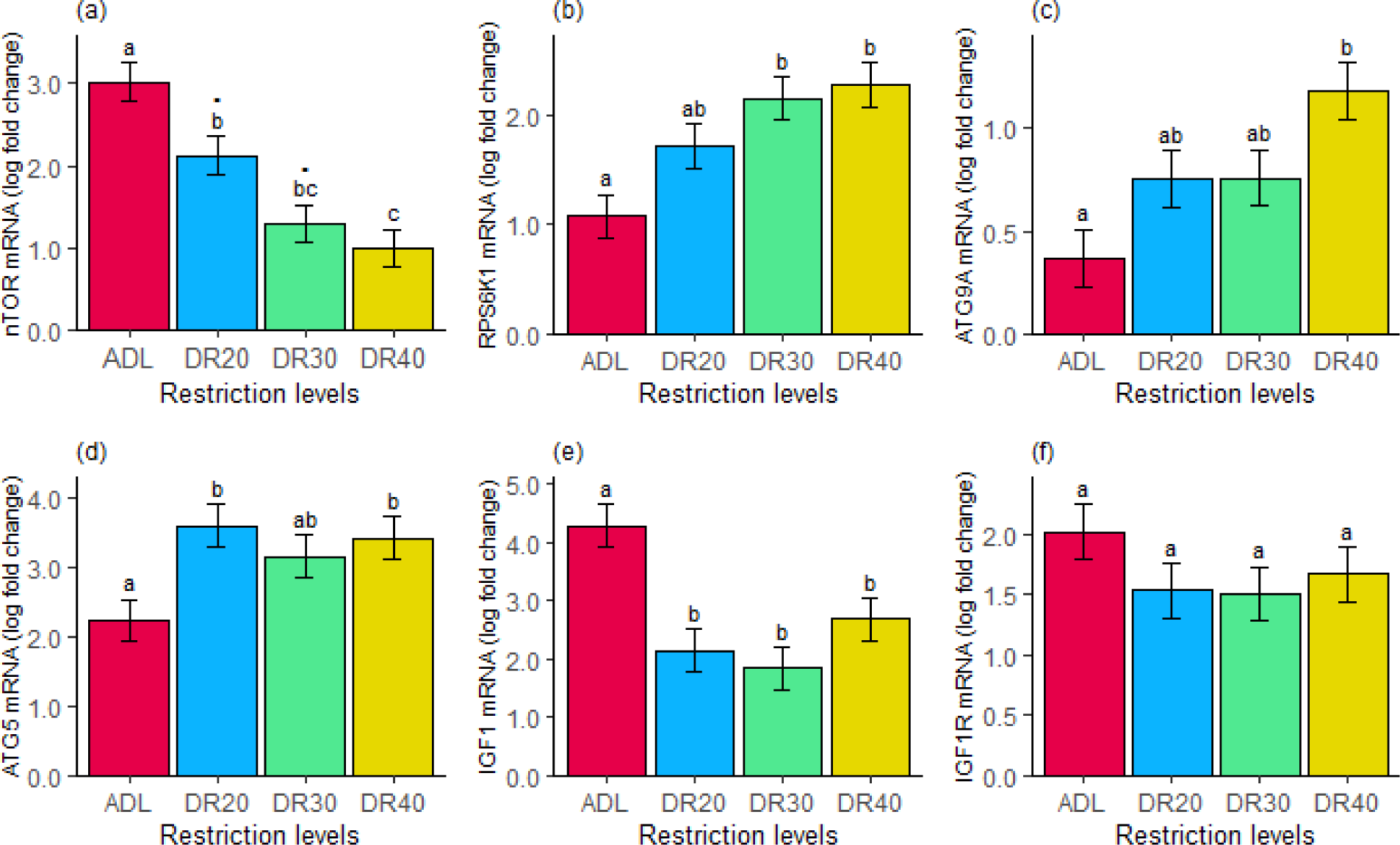
The 14 days dietary restriction showed a significant effect on relative mRNA expression (log of fold change) of (a) mechanistic target of rapamycin (*mTOR*), (b) ribosomal protein S6 kinase 1 (*RPS6K1*), (c) autophagy-related 9A (*ATG9A*), (d) autophagy-related 5 (*ATG5*), (e) insulin-like growth factor 1 (*IGF1*), whereas had no significant effect on (f) insulin-like growth factor 1 receptor (*IGF1R*). Data are represented by the mean ± SE from 8 birds per group. Means followed by a common letter are not significantly different at p < 0.05. Dots on top of letters indicates marginaly significant difference (p<0.1). Abbreviations follow Figure 1.

The DR treatment also significantly decreased *IGF1* gene expression (*F_3,28_* = 8.998, *p* < 0.0002). All the restricted groups had significantly lower *IGF1* gene expression than the ADL-fed controls. Contrary to *mTOR*, the downregulation of *IGF1* did not intensify with the severity of the treatment (Figure 3e, Table S7). Despite a similar trend, *IGF1R* gene expression remained statistically undistinguishable among the four groups (*F_3,28_* = 1.108, *p* = 0.362, Figure 3f). Furthermore, the restriction treatment significantly increased *ATG9A* and *ATG5* gene expression (ATG9A: *F_3,28_* = 5.726, *p* < 0.01, ATG5: *F_3,28_* = 4.117, *p* < 0.05, Figure 3c,d).

### 3.4. Gene expression is related to fitness

The PC analysis indicates that cumulatively 63.7% of the variation is explained by PC1 and PC2 with an eigenvalue greater than 1 (Table S9). PC1 reflects the expression of *ATG9A*, *RPS6K1*, *ATG5*, *mTOR*, and *IGF1*, whereas PC2 reflects mainly the *IGF1R*, *mTOR, ATG9A* and *IGF1* expression (Figure 4). *mTOR* and *IGF1* expressions contributed positively, while *RPS6K1, ATG9A* and *ATG5* expression negatively to variation in PC1 (Table S8). While *mTOR* and *IGF1* are positively correlated, both are negatively correlated with *RPS6K1* and *ATG9A* (Figure S4).

**FIGURE 4.**
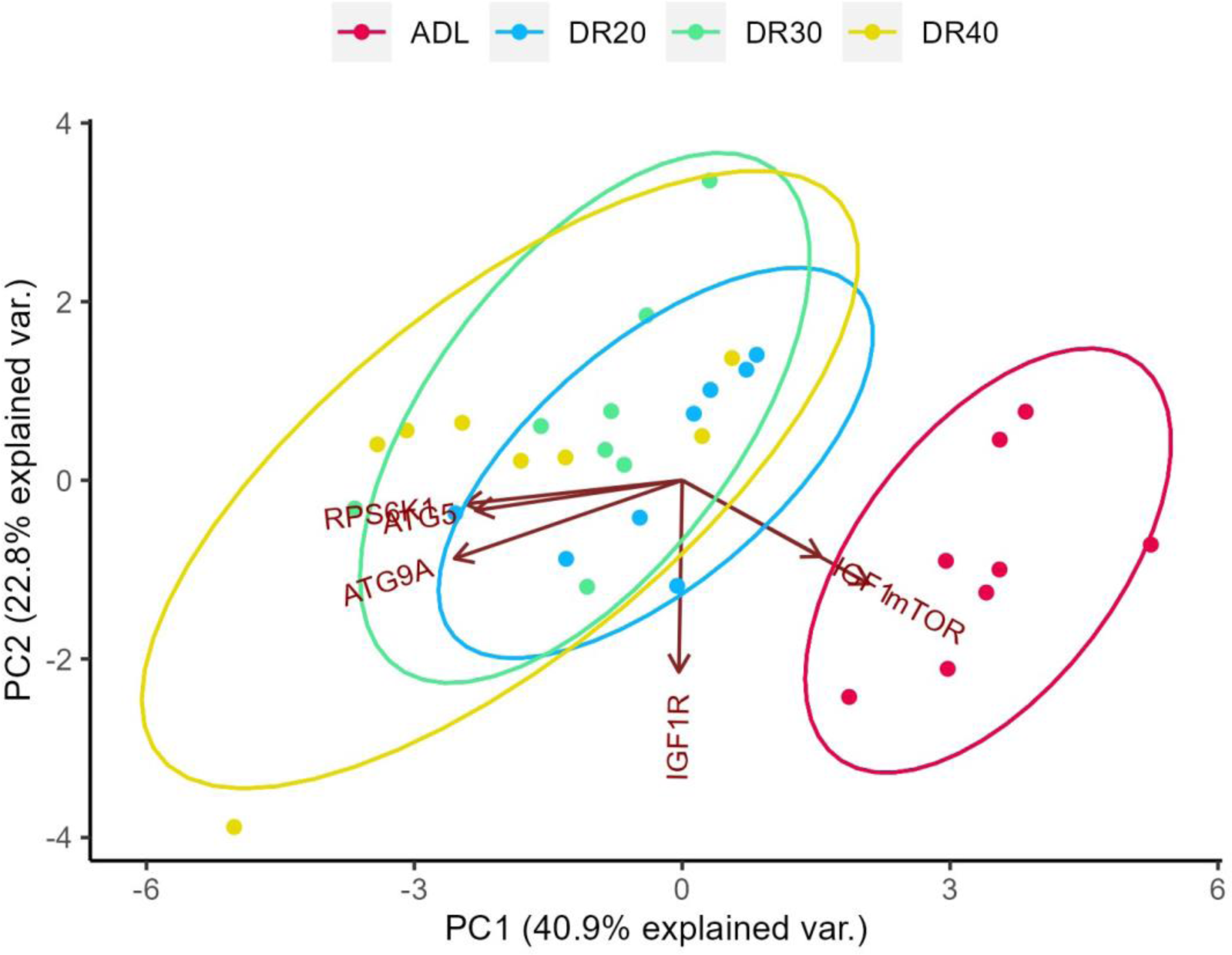
Dimensional indication of PCs of genes expression as independent variables in line with dietary restriction levels clustering. Expression value of *IGF1* and *mTOR* are clustered around ADL group, while catabolic autophagy genes and *RPS6K1* around the restricted groups. Abbreviations follow Figure 1.

Variation in both PC1 and PC2 significantly explained body mass. However, reproductive parameters (egg number and egg mass) were related to only PC1 (Table 2). Therefore, a significant increase in the positive contributor variables of PC1, the *mTOR* and *IGF1*, and a decrease in the negative contributor variables, the *RPS6K1, ATG9A* and *ATG5*, lead to an increase in body mass, egg number and egg mass. A significant decrease in the major contributors to the PC2, *IGF1R* and *mTOR* significantly decrease the body mass.

**TABLE 2.**
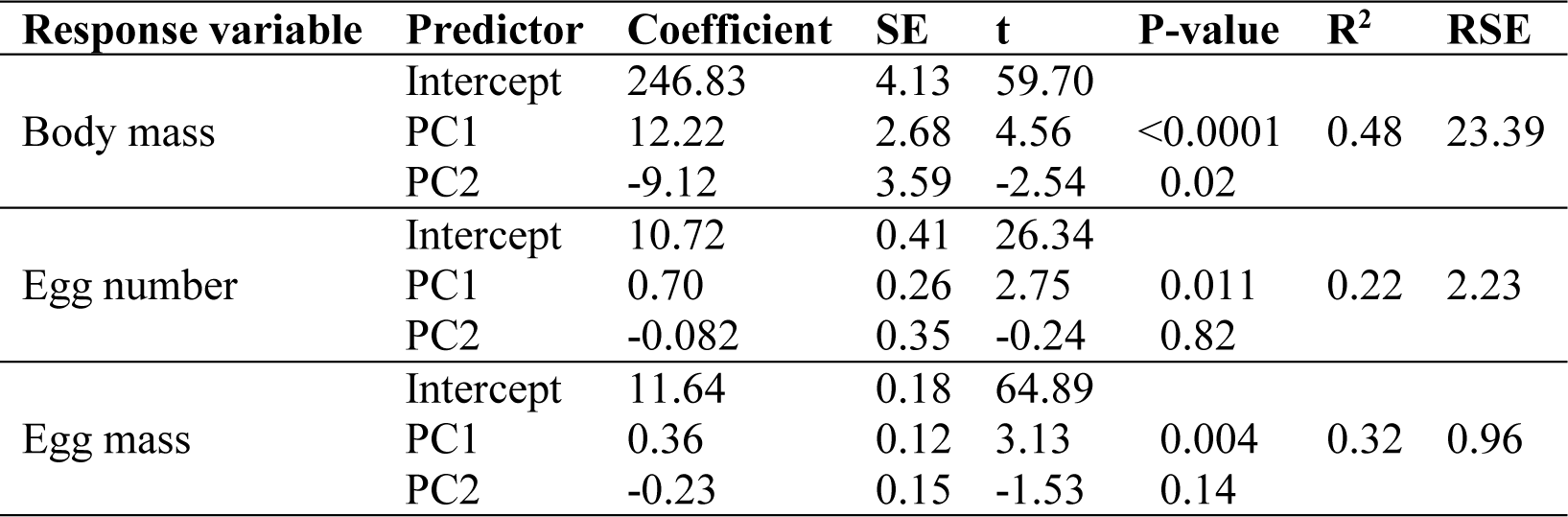
Output of the multiple linear regression of PCs from gene expression predicting body mass, egg number and egg mass. PC1 is positively associated with all fitness components, while PC2 only negatively explained body mass. Principal components are derived from expression value of all genes.

Individually, the expression of mTOR and IGF1 significantly explains all fitness variables positively. The RPS6K1 expression showed a negative relationship with body mass and egg mass, while the ATG9A expression was negatively related to body mass and egg number (Figure S3). As mTOR plays a crucial role in gene transcription, we conducted tests to examine its relationship with the expression of the other genes. Our findings indicate that it significantly accounts for the expression of these genes (Figure S5).

### 3.5. mTOR expression is associated with resource re-allocation

On week 1, only the DR20 group increased relative reproductive investment (p = 0.02), while the ADL controls and the two other treatment groups did not deviate significantly from zero re-allocation, and all treatment groups had similar strategy by the end of the second week (p > 0.4, Figure 5). At the end of the experiment, individual variation in allocation strategy was only related to mTOR expression (t = –3.118, p = 0.004). Irrespective of the treatment, individuals with lower mTOR values were more likely to invest proportionally more in reproduction than individuals with higher mTOR activity (Figure 5).

**FIGURE 5.**
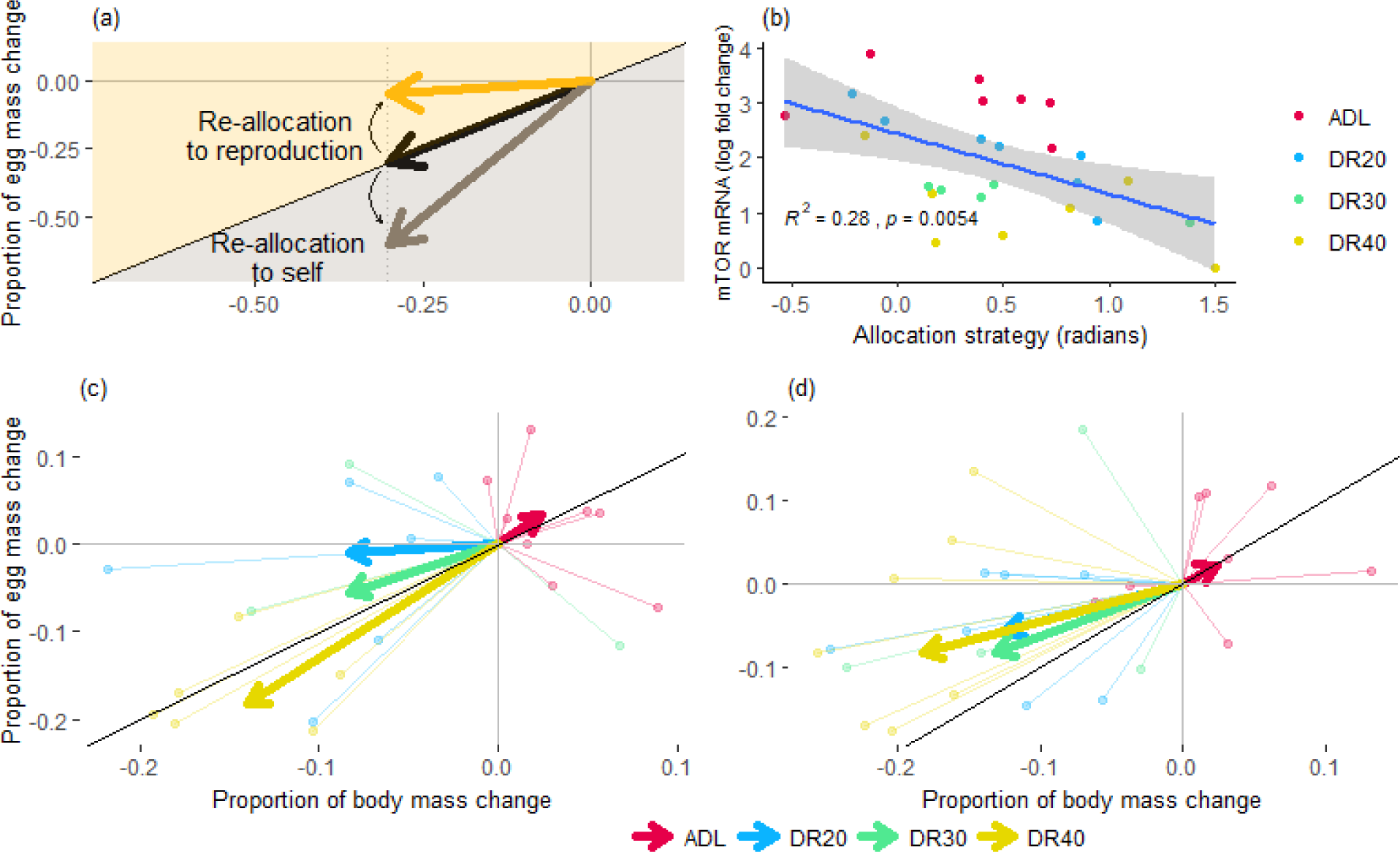
Effect of dietary restriction on resource allocation decision. (a) A conceptual figure illustrating resource allocation decision. The x- and y-axis show the proportional change in body mass and egg-mass, respectively, during the experimental period (compared to the pre-treatment body mass and egg mass, respectively, thus, all vectors start from the origo). The angle of the vectors (radians) illustrates the allocation strategy. The solid black line shows where y = x, i.e. when there is no re-allocation: a unit change in body mass is accompanied by the same amount of correction in egg mass. The space above the identity line indicates reproductive re-allocation: in response to a change of available resources, the individual allocates more to reproduction at the cost of self-maintenance. In contrast, the space under the identity line indicates re-allocation towards self-maintenance because the same change in body mass is associated with a proportionally larger reduction in reproductive investment. (b) Individual allocation strategy is related to mTOR expression. (c-d) The thin lines show individual data points, the thick vectors show the median response of the respective treatment group. In the first week (c), DR20 and DR30 groups invest more in reproduction under a loss of body mass, while DR40 group tends to re-allocate resources to self-maintenance. (d) Over two weeks, all restricted groups show reproductive re-allocation, albeit to different degrees. The longer vectors in the DR40 also illustrate that this treatment imposed a higher cost than in the other two groups (where the lengths of the vectors are similar). The ADL control group remains unchanged over time. Abbreviations follow Figure 1.

## 4. Discussion

Understanding the evolution of resource allocation and its underlying mechanisms remains a major challenge in biology (Moatt et al., 2020; Ng’oma et al., 2017). In this study, we decreased resource availability along a gradient of varying severity and investigated the body mass change, reproductive performance and expression of key genes in the nutrient-sensing IGF-1/mTOR pathway. Our study provided four key results.

First, we found that the severity of the feed restriction affected the body mass differently (Figure 1), indicating that our manipulation of resource availability was successful. Even a mild (20%) feed restriction decreased body mass in the first week, but birds could stabilise their body mass in the second week. The 30% and 40% restrictions resulted in stronger initial body mass losses and birds continued to lose mass in the second week. Birds with higher metabolic rate could be more sensitive to variation in food availability (Brzęk et al., 2012; Zhang et al., 2018).

Second, despite the effective treatments, reproductive performance dropped considerably only when food restriction was more severe. Under 20% food restriction, only the probability of daily egg laying was reduced, indicating that birds were more likely to skip some days in laying, whereas controls reliably laid an egg every day. However, the total number of eggs and the egg mass throughout the study remained similar in DR20 to the ADL-fed birds. In DR30 birds, while the number of eggs remained similar to ADL, the probability of egg laying and the mass of the eggs were reduced, especially in the second half of the experiment. In the most severely restricted hens (DR40), all three reproductive parameters were affected: the probability of egg laying decreased during the study, resulting in fewer and smaller eggs than the controls. However, even under the highest restriction level, the birds continued reproduction, indicating that despite an overall reduction in their resource pool, they managed to keep their reproductive performance, in some cases (DR20) even at the level of the *ADL* controls. However, depending on the magnitude of reduction in the available energy, birds had to face different trade-offs. At a low restriction level (DR20), individuals had to allocate more resources from a limited budget to reproduction, but they could do it without compromising egg size (Figure S2). When resources became more limiting (DR30), birds had to trade-off quality for quantity of reproduction. Under even more challenging conditions (DR40), egg number and mass and also body mass are compromised. Our analysis of resource allocation strategy supports the idea that birds invest in reproduction at moderate restriction (DR20), whereas favour self-maintenance at more severe restriction (DR40) (Figure 5). These results corroborate previous findings (Li et al., 2011; Mahrose et al., 2022), indicating that moderate restriction improves egg production at the expense of egg mass and body mass. Moderate DR has also shown a positive effect on preserving reproductive capacity in mammals (Sun et al., 2021). Mild DR has for years been used in the poultry sector to avoid rapid growth and maintain reproductive life- and health span (Holmes and Ottinger, 2003). A study on rainbow trout indicated that a 20% restriction led to the production of bigger eggs than the ADL fed (Cardona et al., 2019) and suggested that organisms fed ADL seem to invest much of their energy for growth, while the moderately restricted organisms favour investing in their reproductive success.

Third, in response to our treatment, we found characteristic signatures in gene expression patterns. DR downregulated both *mTOR* and *IGF1*, albeit in different ways. The pattern of change in *mTOR* gene expression across treatments groups mirrored the variation in body mass loss and showed a dose-dependent reaction, where the downregulation of *mTOR* gene expression was proportional to body mass loss (Figure 1, 3a, Table S4, S8). In contrast, *IGF1* expression was affected equally under all restricted groups. Nutritional stress has been found to downregulate the expression of the *IGF1* gene, resulting in a decrease in circulating levels of IGF-1. The deficiency of IGF-1 has been observed to have pleiotropic effects (Lodjak and Verhulst, 2020). The result also indicates that the mTOR gene expression is more sensitive to a gradient of nutritional deficiency. Although the specific mechanisms underlying the effect of DR on mTOR gene expression have not been thoroughly investigated, our findings shed light on the similarity between the effect of DR on gene expression and the previously studied effect on the abundance of activated mTORC1 (Velingkaar et al., 2020).

The expression of the mTOR gene is crucial for the cellular production of the mTOR protein, which is then assembled into mTORC1 and mTORC2 complexes along with other component proteins (Szwed et al., 2021). The reduced *mTOR* gene expression could contribute to a lower mTORC1 abundance for activation. Concurrently, DR can downregulate the expression of potential mTORC1 upstream activators. On the other hand, in normal nutritional conditions, activated mTORC1 suppresses mTORC2 activation by phosphorylating RPSK1 (Oh and Jacinto, 2011). Consequently, reduced mTOR gene expression during DR may have a positive impact on the activation of mTORC2 and subsequent cell survival under dietary stress. mTORC2 is also activated by the energy stress sensor AMPK under restriction conditions (Szwed et al., 2021).

Contrary to our assumption, we also found that *RPS6K1* expression increased with the severity of restriction (Figure 3b). In response to phosphorylation by mTOR, RPS6K1 initiates ribosomal translation, consequently promoting cell growth and differentiation (Saxton and Sabatini, 2017). Previous studies have reported reduced *RPS6K1* expression in the liver of overfed geese (Han et al., 2015) and in the brain of restricted mice (Ma et al., 2015). Therefore, we expected that downregulation of *mTOR* would reduce the expression of *RPS6K1* and subsequently reduces body mass (Bae et al., 2012). Despite the predicted decrease in *mTOR* expression, body mass and *RPS6K1* expression were negatively correlated (Figure S3). While a consistent response is expected (Buccitelli and Selbach, 2020), expression of *RPS6K1* gene and the phosphorylated RPS6K1 protein might respond differently to upstream factors. Protein expression of RPS6K1 kinase alone is not sufficient to initiate ribosomal protein translation, rather, it needs to be phosphorylated by the activated mTORC1 kinase (Holz et al., 2005). Hence, the mechanism of action of *mTOR* and *RPS6K1* gene expression, their total protein expression, their phosphorylated expression, their impact on shaping fitness traits are of future research interests. Although there is growing evidence for the correlation between gene expression and protein abundance (Koussounadis et al., 2015; Nie et al., 2006), post-transcriptional modification may alter the biological function of these genes. Currently, the paucity of research reporting the effect of DR on *RPS6K1* gene expression hinders the generalisation of the observed patterns. Nevertheless, our results suggest that *RPS6K1* may be critical in resource allocation decisions. The other genes of interest are the autophagy related genes, *ATG9A* and *ATG5*, the genes involved in autophagosome formation, elongation and closure. Both genes showed a tendency to be upregulated under all restriction groups (Figure 3c).

*mTOR* expression was positively related to *IGF1* expression and its signalling receptor *IGF1R* (Figure S5). The mTOR pathway not only affects translation but is also a key regulator of gene transcription by regulating the activity of specific transcription factors, epigenetic mechanisms or by affecting RNA stability (Laplante and Sabatini, 2013). The modification of transcription factors is important for activation, translocation, interaction, stability and binding (Filtz et al., 2014; Sukumaran et al., 2020). The mTORC1 phosphorylates transcription factors in response to resource availability, which in turn regulate several essential genes. Evidence shows that mTORC1 itself can function as a transcription factor when it is localised in the nucleus (Jiang, 2010; Tsang et al., 2010). Therefore, activated mTORC1 can upregulate the transcription of the *IGF1*, *IGF1R* and the *mTOR* gene itself. In the case of DR, the inhibition of mTOR can lead to a decrease in the expression of genes involved in growth and reproduction, including *IGF1*. Contrarily, under scarce resources (DR), the downregulation of mTORC1 allows the nuclear localisation and activity of Transcription Factor EB (TFEB) and upregulates autophagosome formation through coordinating the expression of genes involved in autophagy such as *ATG9A* and *ATG5* (Martina et al., 2012; Napolitano and Ballabio, 2016). These transcriptional factors are mainly related to maintenance of cellular homeostasis by regulating autophagy and lysosomal genes at the transcriptional level during nutritional deficiency (Inoki et al., 2012; Martina et al., 2012). The correlation analysis in the present study revealed that the expression of *ATG9A* and *ATG5* is negatively related to *mTOR* gene expression (Figure S4), indicating that the downregulation of mTOR mediates the upregulatory effect of DR on autophagy genes.

Finally, we found that variation in the gene expression pattern was coordinated and related to fitness parameters and resource allocation. At a severe DR, the reduced egg number and egg mass are aligned with low *IGF1* and *mTOR* expression, suggesting that these genes are associated with the effect of DR on reproduction (Figure S3). However, individual variation in resource allocation strategy was only related to *mTOR* expression. Stronger restrictions induced an increasing reduction of *mTOR* expression, but irrespective of the treatment, individuals with relatively lower *mTOR* expression had a proportionally larger reproductive investment. This may seem surprising because mTORC1 is required for and thought to promote reproduction (Guo et al., 2018; McLaughlin et al., 2011). The resource re-allocation hypothesis suggests that organisms shift resources between reproduction and somatic maintenance when faced with limited resources (Moatt et al., 2020; Regan et al., 2020), a process mediated by the mTOR pathway. When nutrition is limited, mTORC1 activity is downregulated, triggering alternative pathways (Johnson et al., 2013; Li et al., 2015). In our study, the higher resource re-allocation to reproduction at a lower individual *mTOR* expression may have triggered upregulation of the autophagy pathway and recycling of damaged cell contents as an energy substitution for the nutrient deficit (Adler and Bonduriansky, 2014; Chung and Chung, 2019). The upregulated cellular maintenance helps to preserve the follicle pool and maintain reproductive potential (English and Bonsall, 2019), while activation of autophagy-related genes promotes oocyte maturation (Zhou et al., 2019). Rapamycin treatment, which downregulates mTORC1 was also found to stimulate oocyte maturation by increasing the expression of autophagy-related genes (Lee et al., 2015). In our study, un-inhibition (i.e. upregulation) of recycling mechanisms may have channelled resources towards reproduction. In conclusion, this study revealed that resource limitation-induced allocation trade-offs are associated with a differential expression of nutrient-sensing genes. A limited energy budget induces a lower expression of *mTOR* and *IGF1* and a higher expression of *RPS6K1, ATG9A* and *ATG5* differently at different restriction levels, and leads to overall lower fitness values. However, individuals showing relatively lower *mTOR* expression invest proportionally more in reproduction, which contradicts the established premise that mTOR mediates resource allocation towards reproduction. This apparent paradox may be resolved by a deeper understanding of mTOR’s stimulatory, suppressive and permissive functions.

## Data accessibility statements

All data will be archived in Dryad upon acceptance. R software code for analysis will be archived on relevant database upon acceptance.

## Acknowledgements

We would like to thank James Kachungwa Lugata, Angyal Eszter and Fadella Nur Almira for their help during the experiment. Our thanks also go to Zoltán Németh for valuable discussion and inputs.

## Authors’ contributions

GKR.: conceptualisation, data curation, formal analysis, investigation, visualisation, methodology, writing—original draft, writing—review and editing; SFN.: conceptualisation, methodology, writing—review and editing; BC: data curation, methodology and writing—review and editing; RK: methodology and writing—review and editing; CSz.: methodology, resource and writing— review and editing; GG: methodology and writing—review and editing; LC: conceptualisation, investigation, supervision, funding acquisition, project administration, resources, validation and writing—review and editing; AZL: conceptualisation, data curation, formal analysis, visualisation, methodology, supervision, funding acquisition, project administration, resources, validation and writing—review and editing.

All authors gave final approval for publication and agreed to be held accountable for the work performed therein.

