## Supplementary material for "Dietary restriction and life-history trade-offs: insights into mTOR pathway regulation and reproductive investment in Japanese quails": DR_expression_Supplementary_preprint.pdf

### Expression of genes mediating nutrient sensing pathway explains finesses traits under dietary restriction levels in birds

Gebrehaweria Kidane Reda, Sawadi Fransisco Ndunguru, Brigitta Csernus, Renáta Knop, Csaba Szabó, Czeglédi Levente, Ádám Z. Lendvai

#### Supplemental methods

##### Dietary composition

The standardised basal feed for experimental quails was formulated on a corn, soybean, and wheat basis. They received a breeder quail ration (20% CP; 12.13 MJ/kg ME (National Research Council, 1994)) during acclimation and experiment period (Table S1).

Table S1. Composition and nutrient level of the basal diet

| **Feed ingredients** | **Inclusion rate, %** |
|---|---|
| Corn | 30.37 |
| Wheat | 20 |
| Soybean meal (46% CP) | 34.88 |
| Sunflower oil | 6.79 |
| Limestone | 5.64 |
| MCP | 1.29 |
| Salt | 0.38 |
| DL-Methionine | 0.15 |
| Vitamin and mineral premix[a] | 0.5 |
| **Nutrient content** | |
| Metabolisable energy MJ/kg | 12.13 |
| Crude protein | 20.0 |
| Calcium | 2.5 |
| Available Phosphorus | 0.35 |
| Sodium | 0.15 |
| Methionine | 0.45 |
| Methionine + cysteine | 0.75 |
| Lysine | 1.08 |
| Threonine | 0.74 |
| Leucine | 1.59 |
| Isoleucine | 0.86 |
| Arginine | 1.33 |
| Tryptophan | 0.25 |

[a] 1 kg premix provided: 1000000 NE vitamin A, 200 000 NE vitamin $D_3$, 4900 mg/kg vitamin E, 200 mg vitamin $K_3$, 150 mg vitamin $B_1$, 500 mg vitamin $B_2$, 1200 mg Ca-d-Pantothetane, 400 mg vitamin $B_6$, 2 mg vitamin $B_{12}$, 11 mg biotin, 2502 mg niacin, 60 mg folic acid, 300000 mg choline cloride, 13200 mg Zn, 1920 mg Cu, 9612 mg Fe, 13200 mg Mn, 180 mg I, 42 mg Se, 12 mg Co.

**RNA extraction and cDNA synthesis**

Total RNA from liver tissue was isolated using the TRIzol reagent (Direct-zol™ RNA MiniPrep, Zymo Research Corporation, USA) according to the manufacturer's protocol, including DNA digestion step. Briefly, we lysed 25 - 30 mg liver tissue sample in 600 µl TRIzol Reagent and homogenised it using D1000 handheld homogeniser (Benchmark Scientific, USA). To remove particulate debris from the homogenised tissue, we centrifuged them at 16,000 g for 30 seconds and the supernatant transferred into an RNase-free tube. After we add 600 µl ethanol (95-100%), we mixed it thoroughly and transferred it into Zymo-Spin™ IIC column in a collection tube and centrifuged for 30 seconds again. After adding 400 µl RNA wash buffer and repeating the centrifugation, DNA digestion step was followed by adding 5 µl DNase I (6 U/µl) and 75 µl DNA digestion buffer and incubated at room temperature for 15 minutes to purify the RNA from DNA. After adding 400 µl Direct-zol™ RNA PreWash, we centrifuged for 30 seconds, and repeated this step. For the final wash, we added 700 µl RNA wash buffer and centrifuged for 2 minutes to ensure complete removal of the wash buffer. Finally, we collected the purified RNA by adding 50 µl DNase/RNase-Free water for further RNA quality and quantity check and cDNA synthesis. The RNA concentration in each sample was measured using HTX Synergy Multi-Mode Microplate Reader spectrophotometer at the 260 nm absorbance values (Agilent BioTek, BioTek Instruments Inc, USA), while the purity of each sample was determined by calculating the 260/280 and 260/230 ratios. RNA integrity was checked by 1% agarose gel electrophoresis stained with ethidium bromide. Purified RNA was stored at −80 °C until used for cDNA synthesis on the same day.

Reverse transcription was performed using the qScript cDNA synthesis kit, following the manufacturer's protocol (Quantabio Reagent Technologies, QIAGEN Beverly Inc., USA) in PCRmax Alpha Thermal Cycler (Cole-Parmer Ltd., Vernon Hills, IL, USA). To synthesis the 20 µl final volume cDNA, we used a reaction mix containing qScript cDNA SuperMix (5X reaction buffer containing optimised concentrations of MgCl2, dNTPs, recombinant RNase inhibitor protein, qScript reverse transcriptase, random primers, oligo(dT) primer and stabilisers), 200 ng total RNA and RNase/DNase-free water. The thermal cycling during cDNA synthesis was 25 °C for 5 min (priming), 42 °C for 30 min (reverse transcription) and 85 °C for 5 min (reverse transcriptase inactivation). The cDNA samples were diluted 10-fold and stored at −20 °C for use in Real-time PCR.

**Real-time PCR (qPCR)**

The real-time quantitative PCR was performed using EvaGreen qPCR Mix (*Solis BioDyne*, Teaduspargi, Estonia). The 10 µl final volume of the qPCR reaction mix was contained 5x HOT FIREPol® EvaGreen qPCR mix (*Solis BioDyne*, Teaduspargi, Estonia), 200 nM of each primer, 2 ng cDNA template and distilled water. Intron-spanning gene-specific primer pairs for quails were designed using Oligo7 software and obtained from Integrated DNA Technologies, (BVBA- Leuven, Belgium) (supplementary material, Table S2). We checked for target identity using Primer-Blast software of the National Centre for Biotechnology Information (NCBI) (http://www.ncbi.nlm.nih.gov). The qPCR was performed using the following thermal conditions: 95°C for 12 min (initial activation of the polymerase), 40 cycles of denaturation at 95°C for 15 s, annealing at 60 °C for 20 s and elongation at 72°C for 20 s. At the end of each run amplification specificity of each product was confirmed by melting curve analysis. The qPCR reactions were run in duplicate and the average of each duplicate value was used for subsequent analysis. Amplification and melting curve analysis and monitoring were performed using Agilent AreaMx Real-Time PCR System (Agilent Technologies, USA).

Among the most frequently used reference genes in avian system, beta-actin (*ACTB*), glyceraldehyde-3-phosphate dehydrogenase (*GAPDH*) and 18S ribosomal RNA (*RN18S*), we selected the best reference gene, *ACTB*, using NormFinder, BestKeeper and deltaCt algorithms. Ct values are calibrated against the calibrator sample to calculate the delta-delta Ct value (Azemi et al., 2020; Pabinger et al., 2014). The $2^{-\Delta\Delta Ct}$ method was employed to analyse the relative changes in mRNA expression of target genes (*mTOR*, *RPS6K1*, *IGF1*, *IGF1R*, *ATG9A* and *ATG5*) (Livak and Schmittgen, 2001).

Table S1. Characteristics of the primer pairs used

| Gene* | Primer sequences (5’ → 3’) | Annealing temp (°C) | GenBank Accession No. | Product size (bp) |
|---|---|---|---|---|
| *ACTB*_F | CCC CTG AAC CCC AAA GCC AAC | 56.2 | XM_015876619.1 | 114 |
| *ACTB*_R | ACC AGA GGC ATA CAG GGA CAG C | 56.1 | | |
| *GAPDH*_F | GCA CTG CGC CAC CTT CTC ACT | 57.5 | XM_015873412.2 | 116 |
| *GAPDH*_R | TGA CCA GGC GGC CAA TAC GG | 57.5 | | |
| *RN18S*_F | CCC TGC CGG AGC GTC GAG AA | 59.4 | XR_006936397.1 | 102 |
| *RN18S*_R | CCG GTA ATG ATC CTT CCG CAG GT | 56.1 | | |
| *mTOR*_F | CCG AAG CAT TGA ATT GGC CCT | 53.5 | XM_015882433.2 | 116 |
| *mTOR*_R | CAT CTC TCA AAG GCA GCG GAC C | 55.2 | | |
| *RPS6K1*_F | AGG CAG GAA CCC TCC GTG CAA | 58.8 | XM_015883670.2 | 106 |
| *RPS6K1*_R | AAG CTC AAA CTG CGA AGG GTC GG | 56.9 | | |
| *IGF1*_F | CAC TAT GCG GTG CTG AGC TGG TT | 55.8 | XM_015867574.2 | 118 |
| *IGF1*_R | ATC CCC TTG TGG TGT AAG CGT CT | 55.4 | | |
| *IGF1R*_F | TAC AAC TAC CGC TGC TGG ACC AC | 56.0 | XM_015873184.2 | 107 |
| *IGF1R*_R | AGG CAC TCA GGA TGG CAA CAC | 55.0 | | |
| *ATG9A*_F | CAA CGC CCT CAG GAT CCC CAT | 56.8 | XM_015868966.2 | 69 |
| *ATG9A*_R | ACG ATG CGG GCC TGT ACC TCC | 59 | | |
| *ATG5*_F | ATA GTG GAT TTC GGT ACA TCC CA | 54.03 | XM_015858736.2 | 95 |
| *ATG5*_R | TCC TCC AGA AGC AAT TGG TCG | 55.34 | | |

* *ACTB*: beta-actin, *GAPDH*: glyceraldehyde-3-phosphate dehydrogenase; *RN18s*: 18S ribosomal RNA; *mTOR*: mechanistic target of rapamycin, *RPS6K1*: ribosomal protein S6 kinase 1, *IGF1*: insulin-like growth factor 1, *IGF1R*: insulin-like growth factor 1 receptor; *ATG9A*: autophagy-related gene-9A.

**Supplementary results**

**Effect of dietary restriction on body mass**

Table S3. Summary of the mixed effect model to estimate the effect of dietary restriction on body mass of female and male quails from three time points in two weeks. We fit the model as: body mass ~ treatment × week + (1|block1/birdID), which considers treatment and week as fixed effect and block and individual bird as random effect.

| ***Predictors*** | ***Estimate*** | ***Std. Error*** | ***df*** | ***t-value*** | ***p-value*** |
|---|---|---|---|---|---|
| Intercept | 283.137 | 6.439 | 42.575 | 43.970 | <0.0001 |
| DR20 | -3.450 | 8.660 | 35.460 | -0.398 | 0.6927 |
| DR30 | -12.825 | 8.660 | 35.460 | -1.481 | 0.1475 |
| DR40 | -14.875 | 8.660 | 35.460 | -1.718 | 0.0946 |
| Week 1 | 8.950 | 5.252 | 56.0 | 1.704 | 0.0939 |
| Week 2 | 6.088 | 5.252 | 56.0 | 1.159 | 0.2514 |
| Week 1 * DR20 | -34.638 | 7.428 | 56.0 | 4.663 | <0.0001 |
| Week 1 * DR23 | -32.475 | 7.428 | 56.0 | -4.372 | <0.0001 |
| Week 1 * DR240 | -47.463 | 7.428 | 56.0 | -6.390 | <0.0001 |
| Week 2 * DR20 | -39.288 | 7.428 | 56.0 | -5.289 | <0.0001 |
| Week 1 * DR30 | -43.013 | 7.428 | 56.0 | -5.790 | <0.0001 |
| Week 2 * DR40 | -56.138 | 7.428 | 56.0 | -7.557 | <0.0001 |
| ***Random effects*** | | | | | |
| **Groups** | ***Variance*** | | | **Number of observations** | |
| birdID:block | 189.64 | | | | |
| block | 31.73 | | | 96 | |
| Residual | 110.36 | | | | |

ADL, *ad libitum*; DR20, 20% restriction; DR30, 30% restriction; DR40, 40% restriction; birdID, individual bird; block, experimental block

Table S4. Pairwise comparison of all of the dietary levels based on emmeans for all time points. Randomly allocated groups have no significant difference at the beginning the experiment. Therefore, all differences on the first and second week are due to the dietary treatment. At week 1 (day 7) and week 2 (day 14) all restricted groups showed significantly reduced body mass compared to the ADL group. Additionally, the 40% restricted group also showed significantly reduced body mass compared to the 20% restricted group. On both week 1 and 2 there was significant difference between the 20% and 30% restricted groups, and 30% and 40% restricted groups.

| *Contrast* | *Estimate* | *t-ratio* | *p-value* |
|---|---|---|---|
| **Initial (day 0)** | | | |
| ADL - DR20 | 3.45 | 0.398 | 0.9783 |
| ADL - DR30 | 12.82 | 1.481 | 0.4594 |
| ADL - DR40 | 14.88 | 1.718 | 0.3299 |
| DR20 - DR30 | 9.38 | 1.083 | 0.7023 |
| DR20 - DR40 | 11.43 | 1.319 | 0.5571 |
| DR30 - DR40 | 2.05 | 0.237 | 0.9952 |
| **Week 1 (day 7)** | | | |
| ADL - DR20 | 38.09 | 4.398 | 0.0005 |
| ADL - DR30 | 45.3 | 5.231 | <.0001 |
| ADL - DR40 | 62.34 | 7.198 | <.0001 |
| DR20 - DR30 | 7.21 | 0.833 | 0.8385 |
| DR20 - DR40 | 24.25 | 2.8 | 0.0392 |
| DR30 - DR40 | 17.04 | 1.967 | 0.2194 |
| **Week 2 (day 14)** | | | |
| ADL - DR20 | 42.74 | 4.935 | 0.0001 |
| ADL - DR30 | 55.84 | 5.231 | <.0001 |
| ADL - DR40 | 71.01 | 8.2 | <.0001 |
| DR20 - DR30 | 13.1 | 1.513 | 0.4408 |
| DR20 - DR40 | 28.27 | 3.265 | 0.0124 |
| DR30 - DR40 | 15.18 | 1.752 | 0.3128 |

Standard error = 8.66, degree of freedom = 35.5

Table S5. Pairwise comparison of all of the restriction time points based on emmeans of body mass at each restriction levels. Compare to the initial body mass all restricted groups showed significantly reduced body mass on both week 1 and week 2. The 30% and 40% restricted levels showed significantly and marginal significant body mass reduction from week 1 to week 2, respectively, while the 20% restricted group showed no significant variation from week 1 to week 2 restriction time points. The *ad libitum* group did not show significant variation at all-time points.

| ***Contrast*** | ***Estimate*** | ***t-ratio*** | ***p-value*** |
|---|---|---|---|
| ***Ad libitum* group** | | | |
| Initial – week 1 | -8.95 | -1.704 | 0.2127 |
| Initial – week 2 | -6.09 | -1.159 | 0.4824 |
| Week 1 – week 2 | 2.86 | 0.545 | 0.8495 |
| **20% restricted group** | | | |
| Initial – week 1 | 25.69 | 4.891 | <.0001 |
| Initial – week 2 | 33.2 | 6.321 | <.0001 |
| Week 1 – week 2 | 7.51 | 1.43 | 0.3325 |
| **30% restricted group** | | | |
| Initial – week 1 | 23.52 | 4.479 | 0.0001 |
| Initial – week 2 | 36.92 | 7.03 | <.0001 |
| Week 1 – week 2 | 13.4 | 2.551 | 0.0355 |
| **40% restricted group** | | | |
| Baseline – week 1 | 38.51 | 7.332 | <.0001 |
| Baseline – week 2 | 50.05 | 9.529 | <.0001 |
| Week 1 – week 2 | 11.54 | 2.197 | 0.0805 |

For all groups standard error = 5.25, degree of freedom = 56.0

**Effect of dietary restriction on egg mass**

Table S6. Summary of generalised additive mixed-effects model for effect of dietary restriction on egg mass. We fit the model as: egg mass ~ s(day) + treatment, random = list(birdID=~1), which considers treatment as fixed effect, s(day) as smooth term and individual bird as random effect.

| ***Predictors*** | ***Estimate*** | ***Std. Error*** | ***df*** | ***t-value*** | ***p-value*** |
|---|---|---|---|---|---|
| Intercept | 12.4764 | 0.3116 | 44.429 | 40.044 | <0.0001 |
| DR20 | -0.1832 | 0.4575 | 44.429 | -0.401 | 0.68902 |
| DR30 | -0.9755 | 0.4593 | 44.429 | -2.124 | 0.03442 |
| DR40 | -1.4464 | 0.4435 | 44.429 | -3.262 | 0.00122 |
| ***Approximate significance of smooth terms*** | | | | | |
| | ***df*** | ***F value*** | ***p*** | ***Observation*** | |
| s(day) | 2.202 | 5.441 | 0.00284 | 343 | |

day, restriction days

**Effect of dietary restriction on gene expression**

Table S7. Pairwise comparison for the effect of dietary restriction groups on mRNA expression nutrient sensing mediating genes. All restricted groups showed significantly reduced *mTOR* mRNA expression compared to the ADL group. The DR30 showed a tendency to be significantly lower than the DR20. The DR40 showed significantly lower value than the DR20, whereas there was no significant difference between DR30 and DR40 groups. The DR30 and DR40 groups showed significantly increased *RPS6K1* mRNA expression compared to the ADL group. There was no significant difference among the restricted groups. All restricted groups showed significantly reduced IGF-1 mRNA expression compared to the ADL group. There was no significant difference among the restricted groups. Only the DR40 group showed significantly increased *ATG9A* mRNA expression compared to the ADL group. The DR20 showed significantly increased ATG5 expression than the ADL group. The DR40 also showed marginally significant increase in ATG5 expression.

| *Contrast* | *Estimate* | *t-ratio* | *p-value* |
|---|---|---|---|
| ***mTOR* mRNA expression** | | | |
| ADL - DR20 | 0.898 | 2.733 | 0.0497 |
| ADL - DR30 | 1.730 | 5.264 | 0.0001 |
| ADL - DR40 | 2.032 | 6.182 | <.0001 |
| DR20 - DR30 | 0.832 | 2.531 | 0.0767 |
| DR20 - DR40 | 1.133 | 3.449 | 0.0092 |
| DR30 - DR40 | 0.302 | 0.918 | 0.7957 |
| ***RPS6K1* mRNA expression** | | | |
| ADL - DR20 | -0.642 | -2.294 | 0.1238 |
| ADL - DR30 | -1.078 | -3.850 | 0.0033 |
| ADL - DR40 | -1.203 | -4.293 | 0.0010 |
| DR20 - DR30 | -0.436 | -1.556 | 0.4192 |
| DR20 - DR40 | -0.560 | -2.000 | 0.2122 |
| DR30 - DR40 | -0.124 | -0.444 | 0.9703 |
| **IGF1 mRNA expression** | | | |
| ADL - DR20 | 2.140 | 4.172 | 0.0014 |
| ADL - DR30 | 2.443 | 4.764 | 0.0003 |
| ADL - DR40 | 1.601 | 3.121 | 0.0204 |
| DR20 - DR30 | 0.303 | 0.591 | 0.9339 |
| DR20 - DR40 | -0.539 | -1.052 | 0.7210 |
| DR30 - DR40 | -0.843 | -1.643 | 0.3728 |
| **IGF1R mRNA expression** | | | |
| ADL - DR20 | 0.488 | 1.542 | 0.4267 |
| ADL - DR30 | 0.510 | 1.611 | 0.3887 |
| ADL - DR40 | 0.353 | 1.115 | 0.6836 |
| DR20 - DR30 | 0.022 | 0.069 | 0.9999 |
| DR20 - DR40 | -0.135 | -0.428 | 0.9733 |
| DR30 - DR40 | -0.157 | -0.497 | 0.9592 |
| ***ATG9A* mRNA expression** | | | |
| ADL - DR20 | 0.38234 | -1.960 | 0.2270 |
| ADL - DR30 | -0.38941 | -1.996 | 0.2134 |
| ADL - DR40 | -0.80795 | -4.142 | 0.0015 |

| | | | |
|---|---|---|---|
| DR20 - DR30 | -0.00707 | -0.036 | 1.0000 |
| DR20 - DR40 | -0.42560 | -2.182 | 0.1530 |
| DR30 - DR40 | -0.41854 | -2.146 | 0.1636 |
| ***ATG5* mRNA expression** | | | |
| ADL - DR20 | -1.361 | -3.225 | 0.0159 |
| ADL - DR30 | -0.918 | -2.17 | 0.1547 |
| ADL - DR40 | -1.185 | -2.809 | 0.0420 |
| DR20 - DR30 | 0.443 | 1.049 | 0.7224 |
| DR20 - DR40 | 0.176 | 0.416 | 0.9752 |
| DR30 - DR40 | -0.267 | -0.633 | 0.9206 |

ADL, *ad libitum*; DR20, 20% restriction; DR30, 30% restriction; DR40, 40% restriction; *mTOR*, mechanistic target of rapamycin. *RPS6K1*, ribosomal protein S6 kinase 1; *IGF1*, insulin-like growth factor 1; *ATG9A*, autophagy-related gene 9A; *ATG5*, autophagy-related gene 5

**Principal component analysis of genes against the fitness traits**

Table S8. Contribution of original variables to the principal components. *mTOR* and IGF-1 mRNA expressions positively while *RPS6K1*, *ATG9A* and ATG5 mRNA expression negatively contributed to the PC1. Whereas, *IGF1R, mTOR* and *ATG9A* negatively contribute to the PC2.

| | ***Principal components*** | | | | | |
|---|---|---|---|---|---|---|
| ***Original variables*** | ***PC1*** | ***PC2*** | ***PC3*** | ***PC4*** | ***PC5*** | ***PC6*** |
| *mTOR* | 0.46 | -0.42 | 0.07 | -0.71 | -0.11 | 0.36 |
| *RPS6K1* | -0.49 | -0.09 | -0.26 | -0.21 | 0.79 | 0.14 |
| IGF1 | 0.38 | -0.32 | -0.79 | 0.38 | 0.03 | 0.17 |
| *IGF1R* | -0.01 | -0.78 | 0.30 | 0.26 | -0.03 | -0.51 |
| *ATG9A* | -0.52 | -0.31 | 0.19 | 0.25 | -0.10 | 0.72 |
| *ATG5* | -0.47 | -0.12 | -0.41 | -0.44 | -0.59 | -0.21 |

Table S9. Eigenvalues and respective complained variance of PCs from mRNA expression variables.

| ***PCs*** | ***Eigenvalue*** | ***Individual Variance (%)*** | ***Cumulative variance (%)*** |
|---|---|---|---|
| PC1 | 2.46 | 40.92 | 40.91 |
| PC1 | 1.37 | 22.81 | 63.74 |
| PC3 | 0.86 | 14.26 | 78.00 |
| PC4 | 0.55 | 8.30 | 87.24 |
| PC5 | 0.50 | 7.78 | 95.55 |
| PC5 | 0.27 | 4.46 | 100.00 |

#### Plots

##### Feed restriction has effect on probability of daily egg laying

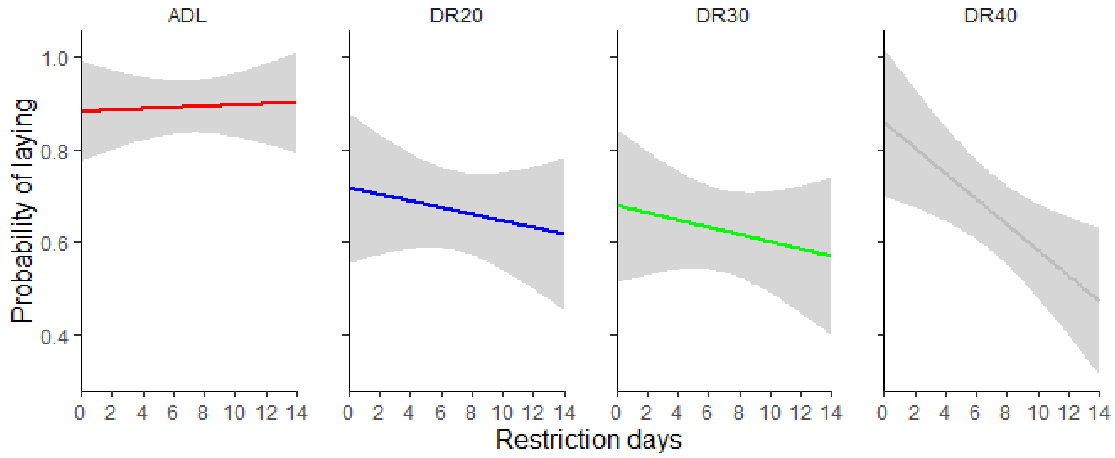


Figure S1. The effect of dietary restriction levels on probability of egg laying. All restriction levels reduced the probability of egg laying. ADL, *ad libitum*; DR20, 20% restriction; DR30, 30% restriction; DR40, 40% restriction.

##### Feed restriction has effect on relative egg mass

At the mid of the experiment period, DR30 and DR40 showed decreasing trend in relative egg mas across the restriction days, while both groups start to reclaim at the last days of the experiment. The DR20 group showed higher relative egg mass than the ADL group across all the restriction days, which indicate that at a moderate DR birds reduced their body mass while keeping egg mass unaffected.

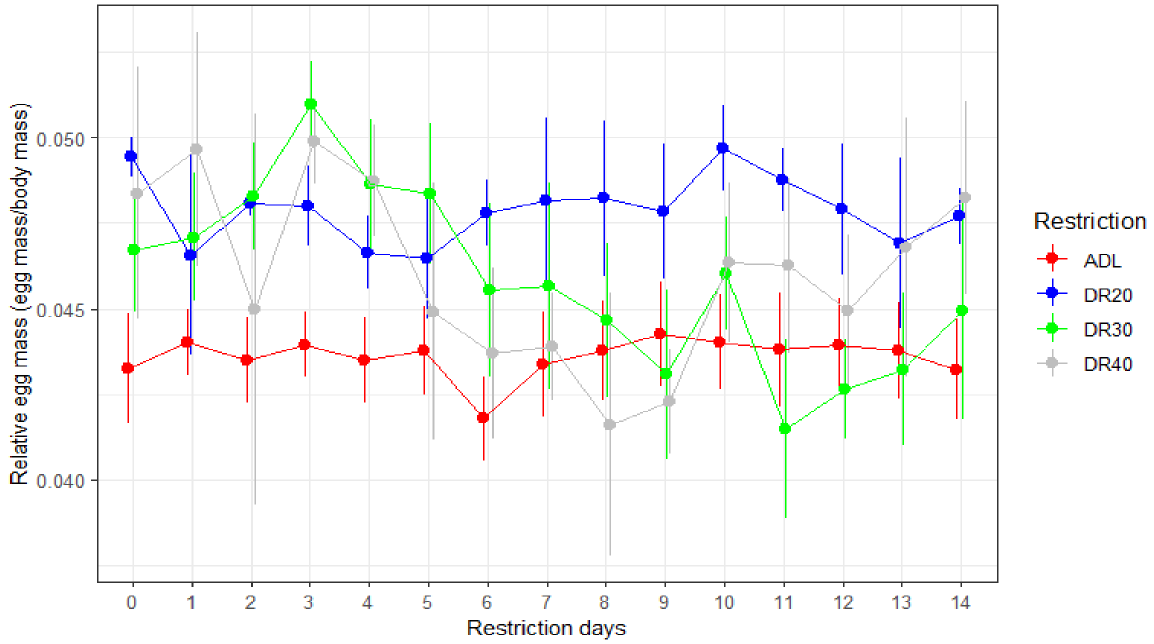


Figure S2. The effect of dietary restriction levels and restriction period on relative egg mass (egg mass divided by average body mass of each birds). ADL, *ad libitum*; DR20, 20% restriction; DR30, 30% restriction; DR40, 40% restriction.

#### Regression of genes mRNA expression variables and fitness traits

We analysed regression of the fitness traits as response variables and the mRNA expression of mediating signalling components as feature variables. The liver *mTOR* relative mRNA expression showed significantly positive relationship with all the three fitness-related traits. Body mass was significantly ($R^2$ = 0.37, $p$ = 0.0004, figure 6a) explained by expression of *mTOR* mRNA. Similarly, the total number of eggs of the 14 days ($R^2$ = 0.34, $p$ = 0.001, figure 6b) and average egg mass of the last 7 days ($R^2$ = 0.18, $p$ = 0.022, figure 6c) were significantly explained by *mTOR* relative mRNA expression (figure 6). There is interaction effect of the *mTOR* relative mRNA expression and DR treatment on the fitness traits.

Similar to *mTOR* relative mRNA expression, the IGF-1 relative mRNA expression also positively explained body mass ($R^2$ = 0.23, $p$ = 0.0058, figure 6d), number of eggs ($R^2$ = 0.34, $p$ = 0.0014, figure 6e) and egg mass of the last 7 days ($R^2$ = 0.13, $p < 0.05$, figure 6f). The *RPS6K1* relative mRNA expression showed significantly negative relationship with body mass ($R^2$ = 0.27, $p$ = 0.0025, figure 6g) and egg mass of the last 7 days ($R^2$ = 0.25, $p < 0.0058$, figure 6i), while it has no significant relationship to the number of eggs ($R^2$ = 0.023, $p$ = 0.42, figure 6h). The *ATG9A* relative mRNA expression negatively explained the body mass ($R^2$ = 0.17, $p$ = 0.025, figure 6j) and egg number ($R^2$ = 0.19, $p$ = 0.02, figure 6k), while no significant relationship with egg mass of the last 7 days ($R^2$ = 0.083, $p$ = 0.13, figure 6l).

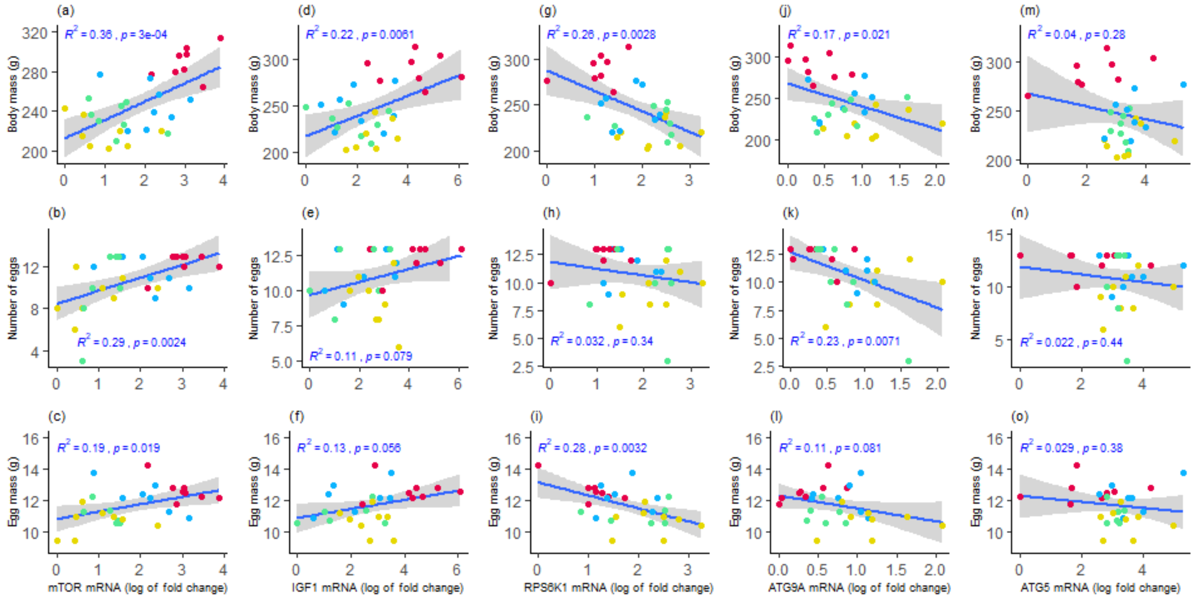


Figure S3. Linear regression analysis between liver relative mRNA expression (log of fold change) with fitness traits. *mTOR* relative mRNA expression positively explains (a) body mass, (b) total egg number and (c) egg mass. IGF-1 relative mRNA expression positively explains (d) body mass, (e) total number of eggs and (f) egg mass. *RPS6K1* relative mRNA expression negatively explains (g) body mass and (i) egg mass, and no significant relationship with (h) total egg number. *ATG9A* relative mRNA expression negatively explains (j) body mass and (k) and egg number, and (l) no significant relationship with egg mass. Egg mass was the mean egg mass of the last 7 days of the experiment. Colours representation, red, *ad libitum*; blue, 20% restriction; green, 30% restriction; black, 40% restriction.

**Pearson correlation of nutrient sensing pathway mediating genes**

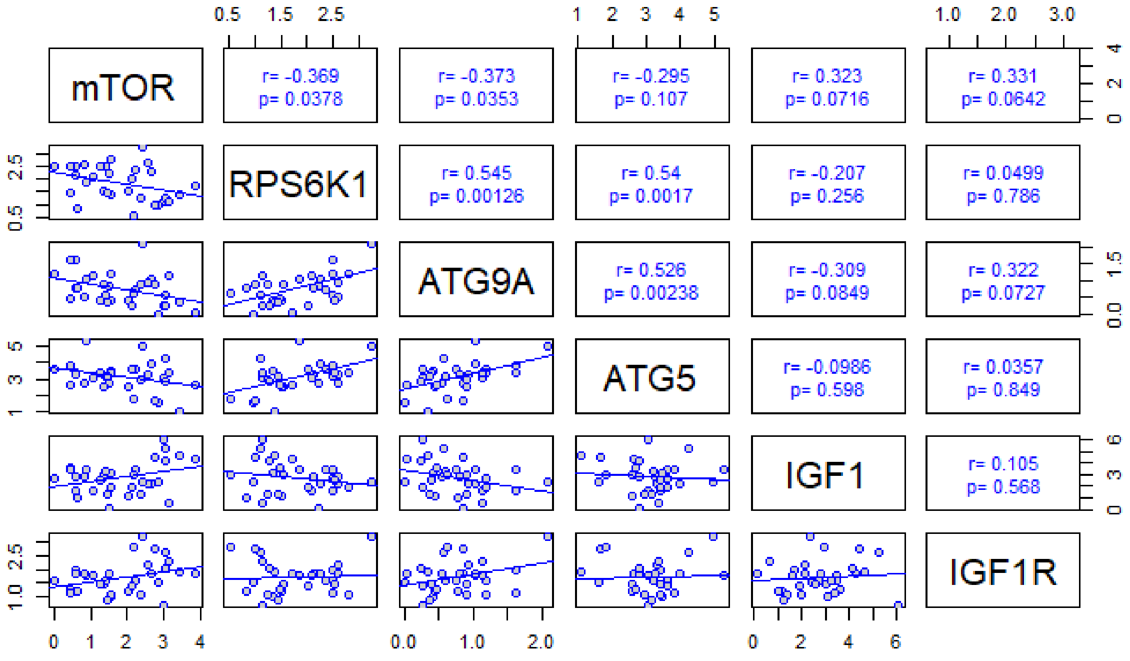


Figure S4. Pearson correlation among mRNA expression of genes mediating nutrient sensing pathway. *mTOR* showed significantly positive correlation with IGF-1 and negative correlation with *RPS6K1*, *ATG9A* and ATG5 expression. RPS6K1, ATG9A and ATG5 showed positive correlation to each other. IGF-1 expression showed negatively marginal significant correlation with *ATG9A* expression.

**_mTOR_ explains expression of other mediating genes**

The *mTOR* relative mRNA expression marginally explains IGF-1 ($R^2$ = 0.11, $p$ = 0.077, figure S4a) and *IGF1R* ($R^2$ = 0.11, $p$ = 0.073, figure S4b) relative mRNA expression. Unexpectedly, the *RPS6K1* relative mRNA expression showed significantly ($R^2$ = 0.14, $p$ = 0.047, figure S4c) negative relationship with *mTOR* mRNA expression. The autophagosome formation gene, *ATG9A* mRNA expression showed significantly ($R^2$ = 0.16, $p$ = 0.032, figure S4d) reduced value of relative mRNA expression with increasing value of *mTOR* mRNA expression.

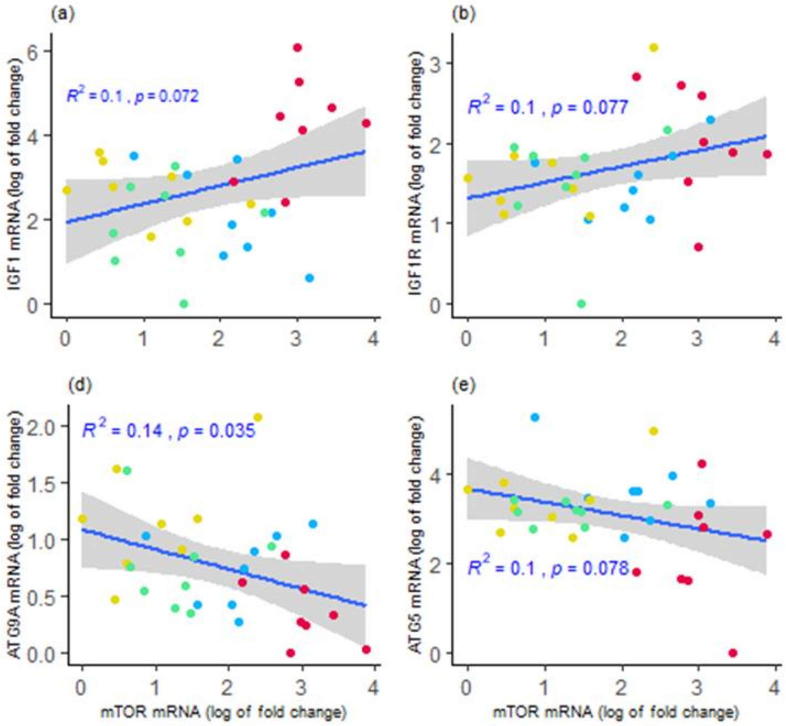


**Figure S5**. Linear regression of liver *mTOR* relative mRNA expression (log of fold change) with mRNA expression of other mediating genes. *mTOR* relative mRNA expression showed a tendency to positively explain IGF-1 and IGF-1R relative mRNA expression (a, b); *mTOR* relative mRNA expression significantly negatively explain *RPS6K1* and *ATG9A* relative mRNA expression (c, d); mTOR has no significantly explain ATG5 (e). Colours representation, red, *ad libitum*; blue, 20% restriction; green, 30% restriction; black, 40% restriction.
